# Neural Representation Precision of Distance Predicts Arithmetic Performance

**DOI:** 10.1101/2024.08.01.605970

**Authors:** Hui Zhao, Wang Qi, Jiahua Xu, Yaxin Yao, Jianing Lv, Jiaxin Yang, Shaozheng Qin

## Abstract

Focusing on the distance between magnitudes as the start point to investigate the mechanism of relationship detecting and its contribution to mathematical thinking, this study explores the precision of neural representations of numerical distance and their impact on arithmetic performance. By employing neural decoding techniques and representational similarity analysis, the present study investigates how accurately the brain represents numerical distances and how this precision relates to arithmetic skills. Thirty-two children participated, completing a dot number comparison task during fMRI scanning and an arithmetic fluency test. Results indicated that neural activation patterns in the intra-parietal sulcus decoded the distance between the presented pair of dots, and higher precision in neural distance representation correlates with better arithmetic performance. These findings suggest that the accuracy of neural decoding can serve as an index of the neural representation precision and that the ability to precisely encode numerical distances in the brain is a key factor in mathematical abilities. This provides new insights into the neural basis of mathematical cognition and learning.

**Highlights:**

1. Utilizing representational similarity analysis and neural decoding techniques, the research proposes that the accuracy of neural decoding serve as an index of neural representation precision.
2. The precision of neural representation of numerical distances in the brain predicts task performance and math arithmetic proficiency in children.
3. These findings imply the potential significance of relational information in general cognition and learning beyond mathematical learning.

## 1. Introduction

Figuring out the relationship between items is fundamental for individuals to acknowledge the world. People create travel itineraries by labeling the building around as close or far for orientation, compare similarities and differences between different words to build semantic concepts, and assess familiarity and strangeness with others to navigate social interactions. Mathematics, to some extent, is a scientific discipline that accurately describes relationships and rules between objects using quantitative language. These relationships can be described creatively and universally, e.g., C^2^ as the relationship between E and m. However, determining the difference between magnitudes and becoming aware of the distances between them is perhaps the simplest and most basic form of relationship detection.

The capacity to distinguish different magnitudes is considered intuitive and foundational to mathematical cognition (Ansari, 2008; Brannon, 2006; Dehaene, 1992). However, there has not been a study focusing on the representation of distance between magnitudes, let alone its representation precision. Moyer & Landauer (1967) found that the participants reacted faster and more accurate when the magnitude distance between two numbers was larger, this so-called distance effect has been extensively replicated in numerical magnitude comparison research. The distance effect follows Weber’s Law, with reaction time and accuracy varying as a function of distance/ratio between the numerical sets (Hais, Wazny, Toomarian, & Adamo, 2015). However, the distance effect per se cannot indicate the precision of distance representation, nor can it determine the precision of magnitude representation. On one hand, a greater distance effect might imply the capacity to detect the different distances between magnitudes, suggesting a clear or a possible accurate representation of magnitude. That is why the ratio/distance effect was increased over development (Haist et al., 2015; Lyons, Nuerk, & Ansari, 2015). On the other hand, subjects might react faster even with very close number sets when they have more accurate magnitude/distance representation, which would result in decreasing distance effect. This could explain why some researchers also found a decreasing distance effect over development (Holloway & Ansari, 2010), and further correlating with higher numerical acuity and even better math achievement (Ashkenazi, Mark-Zigdon, & Henik, 2009; De Smedt, Verschaffel, & Ghesquïre, 2009; Mundy & Gilmore, 2009). In contrast to the paradox of the distance effect, the Weber fraction value is well-accepted to detect the acuity of distinguishing two magnitudes/numbers (Pica, Lemer, Izard, & Dehaene, 2004). Halberda, Mazzocco, and Feigenson (2008) further claimed that the numerical acuity was even correlated with math achievement. They found that the Weber fraction of 14-year-old children correlated with their past performance on standardized math achievement tests. And those results were repeated by Libertus (2011) and even across life span (Halberda et al., 2012; Libertus, Brannon, et al., 2011). However, it remains unclear whether the acuity in magnitude comparison tasks derives from precise distance representation.

No brain imaging study to date has illustrated distance representation in number processing or math learning research as well. However, distance coding, as revealed through multivariate voxel pattern analysis (MVPA) and neural representational similarity analysis(RSA), has been considered in the research on navigation in physical or cognitive maps, underscoring the importance of distance representation between the positions is crucial for even cognition or learning (Theves, Fernandez, & Doeller, 2019; Viganò & Piazza, 2020). These studies found involvement of the hippocampus or entorhinal cortex in distance decoding. Similar method has been used in numerical-related studies to address magnitude representation (Dormal, Andres, Dormal, & Pesenti, 2010; Eger, Pinel, Dehaene, & Kleinschmidt, 2015). These studies primarily focus on the parietal area, as it is extensively reported in numerical-related and math-related research, yet very few have addressed the precision of magnitude presentation. Zhang et al. (2023)have claimed that the summed function connectivity of numerosity network could serve as a bio-marker for non-verbal number acuity, but this functional connectivity is merely associated with acuity and does not directly quantify the degree of acuity. Some researchers have raised the issue of neural fidelity employing neural pattern similarity in episodic memory or stimulus representation tasks (Rothlein, DeGutis, & Esterman, 2018; Zheng et al., 2018). However, precise representation requires not only mean a stable representation, as indicated by similar neural pattern induced by repeated representation of same stimulus, but also the clear distinction between different stimuli. Recently, Barretto-García et al. (2023) used the slope of the psychometric curve for a probability distribution to select a given magnitude in the magnitude comparison task based on the noisy logarithmic coding (NLC) model (Khaw, Li, & Woodford, 2021). They found out that the magnitude neural representation acuity was related to risk decision. However, determining the receptive field of number is challenging when trying to define a concrete receptive field for an abstract distance, and no paper has yet revealed the precision of distance presentation.

The updated neural decoding method offers an insightful methodology to investigate both distance and magnitude representation precision. The machine learning model trained to recognize the patterns of neural activity of a given numerical distance/magnitude and then be able to distinguish the neural response to different distances/magnitudes. When the representation of the target is stable and unique, decoding performance would be ideal, with higher decoding accuracy indicating more accurate neural representation. Previous research has shown that people can decode numerical magnitude in both non-symbolic and symbolic form, visually or auditorily (Bulthé, De Smedt, & Op de Beeck, 2015; Damarla, Cherkassky, & Just, 2016; (Lyons, Ansari, & Beilock, 2015). However, no research has yet used this method to represent acuity and distance decoding.

So the present study would investigate the characteristics of precise neural distance representation and its contribution to magnitude comparison task performance and arithmetic capacity. First, the effect of distance representation and magnitude representation will be tested using the neural tuning curve model (Lyons, Ansari, et al., 2015; Merten & Nieder, 2009) during a magnitude comparison task. Then, the precision of distance representation will be tested by means of neural decoding, and its correlations with task performance and arithmetic performance will be examined.

## 2. Materials and Methods

### 2.1 Participants

A total of 32 healthy children were initially recruited for this study. Three participants were excluded due to either incomplete imaging data (N = 1) or the presence of structural image anomalies (N=2). The ART toolbox was employed to detect artifact points with two criteria: (1) frame to frame displacement > 0.5 * voxel size; (2) signal amplitude surpassing three standard deviations above the average whole-brain signal. Subjects were eliminated when more than one-third of the time points exhibited artifacts. Ultimately no subject was eliminated due to measurement artifacts. Consequently, 29 participants (16 males, mean age = 8.61 ± 0.92) were included in the analyses. All participants were right-handed, had normal or corrected to normal vision, and no history of neurological diseases. Written informed consent was obtained from the parents of all child participants, and the children themselves also gave their assent to participate in the study. Ethical approval of the study was granted by the Institutional Review Board of the Institute.

### 2.2 Stimuli and Experimental Design

Each participant completed the dot number comparison task in the scanner and the basic arithmetic fluency test outside the scanner.

During the fMRI scanning participants were centrally presented with arrays of dots successively. Participants were instructed to respond with one of the two buttons depending on whether the first or second number was larger, if the first number was larger then press the leftmost button with the thumb of left hand, if the second number was larger then press rightmost one with the thumb of right hand, button pressed was balanced across trials. The magnitude of the two dot patterns and the distance in-between are orthogonalized. Participants performed 3 experimental blocks, each block containing 24 trials. The details were shown in Fig 1.

**Figure 1.**
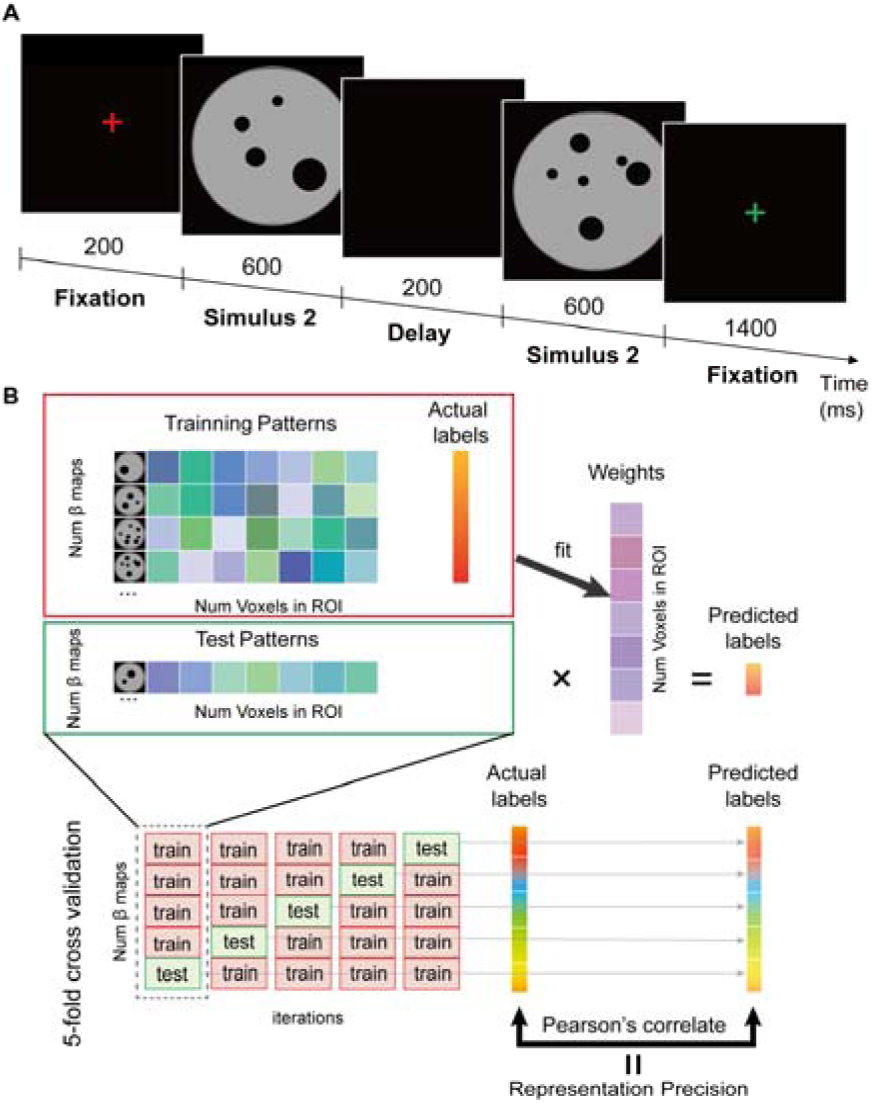
Illustration of The Task Design and Decoding Procedure. Note: (A) The illustration of the dot pattern comparison procedure. Subjects need to compare the two numerical stimuli and decide which one was larger. (B) We first divided the data into a training set with 4/5 trials, and a test set with the remaining 1/5 trials. In the training phase, the pattern from 4/5 training trials and their actual labels (distance or magnitude) were used to fit a model (set of weights on each voxel). During the testing phase, the test patterns were multiplied by the weight vector to generate predicted labels (which cloud take on any real value). A 5-fold cross validation was conducted which iteratively left out 1/5 trials. Performance was assessed by computing the correlation between the actual and predicted labels (aggregated across iterations).

### 2.3 Weber Fraction Estimation

The Weber fraction estimated each participant’s minimum distinguishable difference between the number of dots and which was also known as the ANS (Approximate number system) acuity. Weber Fraction score was accessed for 29 subjects using the ACC of dot comparison task. The percentage correct was modelled for each individual subject as 1-error rate (Halberda et al., 2008), where error rate is defined as:

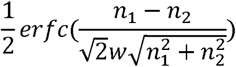

Where *erfc*(*x*) is the complementary error function, which is associated with the integral of the normalized Gaussian distribution. This model describes the percentage of correct response as a function of the Gaussian approximation of number representations for the two dot sets presented in each trial, namely *n_1_* and *n_2_* (i.e., the first and second sets of dots), with a single adjustable parameter, the Weber fraction (*w*).

### 2.4 Arithmetic Fluency Test

The basic arithmetic fluency test assessed the subject’s math performance by performing tasks including addition, subtraction, multiplication and division [adapted from (Wei et al., 2012)]. A math problem was presented at the center of the screen, with two answer choices displayed below it. The subject press a key to select the correct answer. Each task had an overall time limit of 50 seconds. The program recorded the subject’s accuracy and reaction time for each problem. The test included four types of problems: addition, subtraction, multiplication and division. The subject’s performance on each type of problem was compared with the dataset of primary school students in the city(Li, Yang, Lu, Wang, & Zhao, 2015). The subject’s arithmetic performance index (***Z_arithmetic***) was calculated as follows:

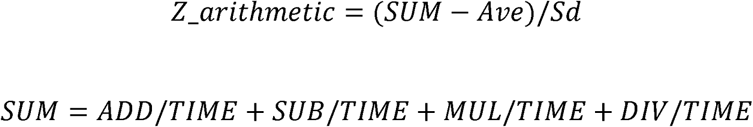

Where, ***ADD***, ***SUB***, ***MUL*** and ***DIV*** are respectively the difference between the number of correct and incorrect problems for each condition. ***TIME*** is the time taken to complete each type of problem. ***Ave*** and ***Sd*** is the average and standard deviation of the group of primary school students for each grade(Supplementary Table 1).

### 2.5 MRI Acquisition Procedure

Images were acquired on a 3.0 T Siemens MRI scanner using a 64 channel head coil. Visual stimuli were projected onto a screen behind the scanner, which was made visible to the participant through a mirror attached to the head coil. Functional images were collected using a single-shot echo-planar imaging (EPI) sequence (axial slices, 33; slice thickness, 3.5mm; gap, 0.7mm; TR, 2000ms; TE, 30ms; flip angle 90°; voxel size, 3.125×3.125×3.125 mm^3^, FOV, 200×200mm2; volume, 158), while structural images were acquired through three-dimensional sagittal T1-weighted magnetization-prepared rapid gradient echo (144 slices; slice thickness, 1.33mm; TR, 2530ms; TE, 3.39ms; voxel size, 1.0×1.0×1.33mm3, flip angle, 7°; FOV, 256×256mm).

### 2.6 Preprocessing and Modeling of Neuroimaging Data

Task-fMRI data were preprocessed and analyzed for each participant using Statistical Parametric Mapping (SPM12; http://www.fil.ion.ucl.ac.uk/spm). For each participant, the first 5 volumes (10 s) of each run were discarded for signal equilibrium. The preprocessing of functional images included slice timing correction and head motion correction. Subsequently, T1-weighted image were co-registered to each participant’s functional image, then segmented into gray matter image and spatially normalized into a common stereotactic Montreal Neurological Institute (MNI) space and resampled into 3-mm isotropic voxels.

The normalized, smoothed EPI volumes were entered into a GLM to generate beta maps as model input for decoding analysis, with second stimulus in each trial modeled as a condition. To ensure that the population neural patterns were not affected by confounding factors such as distraction or random choice, only correct trials were modeled, resulting in an average of 64.8 correct conditions out of 72 total trials, along with 1 error condition (treating all error trials as a single condition) for each participant. The onsets of these conditions were convolved with a standard hemodynamic response function. Additionally, 24 regressors of no interest corresponding to the 24 motion parameters, spike regressors detected using the ART toolbox, a constant regressor for each run, and a high-pass filter with a cut-off set at 90 s were included in the GLM. The resulting beta estimates for the “second number” stimulus conditions were z-scored within each session and then employed in the multi-voxel decoding analysis.

Prior to conducting RSA analysis, the unsmoothed EPI volumes were entered into the GLM for all 72 trials. Each trial was treated as an individual regressor in the GLM. Furthermore, regressors representing the first stimuli condition and the rest condition were established. Spike regressors, identified using the ART toolbox, a constant regressor for each run, and a high-pass filter with a cut-off frequency of 90 seconds were included as nuisance regressors to remove any unwanted variance. Subsequently, t maps were derived from each regressor to compute the dissimilarity matrix.

### 2.7 ROI Definition

Bilateral Intra-parietal sulcus(IPS), Left IPS and Right IPS were selected as regions of interests (ROIs) using SPM anatomy toolbox (Eickhoff et al., 2005) in the analysis of magnitude and distance representation (Fig3.A). Additionally, hippocampus (HIPP) from the WFU pickAtlas (Maldjian et al., 2003) was selected as the contrast for the analysis of distance representation, generating Bilateral HIPP, Left HIPP and Right HIPP as ROIs (Supplementary. Fig2.A).

### 2.8 Decoding Analysis

Linear Support Vector Regression (SVR) with a regularization parameter C = 1 was used in scikit-learn (http://scikit-learn.org/stable/) to decode number magnitude or distance of each trial. A 5-fold cross-validation method was utilized, and the decoding performance was evaluated by calculating Pearson’ r between target labels and predictions across entire cross-validation circle. Here, images were smoothed by an isotropic three-dimensional Gaussian kernel with 6 mm full-width at half-maximum during preprocessing step. Note that, in the magnitude decoding, to mitigate the mixed decoding of hand response, we split the trials into two sets of “1-4” (also always left handed response) and “6-9” (right-hand). We then performed decoding within each of these sets separately and combined the predictions to calculate the overall decoding performance. A Fisher-Z transformation was conducted on Pearson’s *r*, defining *z_r_* as the index of representational precision (RP), which measured how precisely neural population activity represents number distances.

Decoding performance was tested on group level using one sample t-tests and permutation test. In the permutation test, the labels of each trial were shuffled, and 1000 iterations of the entire decoding procedure were performed to obtain the null distribution of decoding performance, and *p* value was calculate by the number of times that the decoding performance in the set of 10,00 permutation tests outperformed the real decoding performance divided by 1000. P values were corrected for multiple comparisons using FDR (Benjamini & Hochberg, 1995).

### 2.9 Representational similarity analyses

#### 2.9.1 Neural Representational Dissimilarity Matrices (RDMs)

The Neural representational dissimilarity matrices(RDMs) was calculated using T map for each trial generated from GLM(Misaki et al., 2010). The Dissimilarity between two trials were computed as 1-r, where r was calculated using partial Pearson correlation including baseline activity as a covariate. This serving as a statistical means to reduce the influence of non-interest elements, such as shared vascular, neural, and imaging factors which could lead to high correlations unrelated to the functional elements of interest (Lyons, Ansari, et al., 2015).

#### 2.9.2 Behavioral Representational Dissimilarity Matrix

For behavioral data, the Euclidean distance of response times (RT) between pairs of trials was computed to generate a trial-by-trial behavioral RDM with dimensions of 72 by 72.

#### 2.9.3 Computation of Distance and Magnitude Tuning Curve Models

Magnitude model used simulated neuronal tuning curves for number magnitude, where the tuning curve defined as a Gaussian function with widths linearly increasing with number magnitude. The hypothesis posits that the observed neural patterns of similarity relations between pairs of numbers correspond to the degree of overlap predicted by these tuning curves, as validated by Lyons et al. (2015). In the present study, the dissimilarity of magnitude was defined as the 1 - overlap (calculated by the proportion of those curves that overlap with one another) of two numbers. By assessing the overlap of curves for any two magnitudes, the magnitude tuning curve overlap matrix was readily obtained (Fig2A, D). To generate the trial-by-trial magnitude representational dissimilarity matrices (RDMs), the magnitude of the second stimulus in each trial was used as a category label. Each value in the RDM was then filled with the overlap value from the curve overlap matrix corresponding to the magnitudes of the respective trials (Fig4A). This means that the dissimilarity between any two trials is quantified by how much their corresponding magnitude tuning curves overlap.

The distance RDM was constructed based on the hypothesis that distance tuning functions similarly to the magnitude tuning curve (Fig2C), with the only difference that the x-axis represents distance between to numbers (ranging from 1 to 6 in our experiment). To examine this hypothesis, these curves were employed to generate trial-by-trial distance RDMs. Importantly, distance RDMs were utilized in both behavioral and neural representational similarity analyses (RSA).

Overlap RDM was utilized in the behavioral RSA analysis. The overlap of pairs of numbers within a single trial was defined as an "overlap curve", representing the trial’s response evoked by those two numeric magnitudes. Since each trial involves two numbers (unlike fMRI data that can be separated), across-trial similarity was computed by measuring the overlap of the overlap curve with other trials, referred to as the "overlap of overlap" (Fig2B). In the experiment, which comprised a total of 12 pairs of overlap categories (Fig2E), trial-by-trial overlap RDM (Fig4A) were derived, similar to magnitude RDMs. For clarity, here we use the term "similarity," but the actual computation is dissimilarity, consistent with our previous analyses.

In order to control potential confounding factors, we also constructed 2 confounding model that characterizes low-level visual similarity and response similarity. (1) Visual RDM: To control for the low-level visual similarity effects between dot images, we calculated the pixel dissimilarity of image pairs. This dissimilarity was computed by Pearson’s correlation distance between the grayscale values of images. (2) Response RDM: A button-press RDM was constructed to control for the effects of motor responses in the dot comparison task. This was computed as the absolute difference between the button press responses (1 for left index finger; 0 for right index finger) recorded during scanning.

#### 2.9.4 Compare RDMs

The neural RDM and behavioral RDM were then compared with model-based RDMs using Spearman’s rank correlation to obtain the “raw effects” of each computation-model RDM. To account for potential confounding factors, we further applied partial Spearman’s rank correlation, controlling for low-level factors (visual RDM and response RDM) to obtain the “preliminary effects” of each computation-model RDM. Subsequently, we controlled for another computation models (e.g., controlling magnitude model when testing distance model) when tested one model to isolate the “unique effect” of each model. The resulting *r* values were fisher-z transformed to conduct group-level one-sample t-test to test whether the RSA results of the theoretical models were significantly above zero. Multiple comparisons were corrected using FDR method (Benjamini & Hochberg, 1995).

### 2.10 Correlation analysis

The decoding accuracy was correlated with the task and arithmetic performance to reveal the importance of neural representational precision.

Specifically, partial Pearson correlations between Representational Precision (RP, including distance RP and magnitude RP) and math performance was tested. Reaction Time (RT) and accuracy (ACC) in the dot comparison task was controlled for to ensure that any observed correlation was not attributable to behavioral performance. When testing the correlations between RP and Weber fraction (w), we only controlled for RT, as w was derived from ACC, which could introduce spuriousness or contamination (Becker et al., 2016),

The significance level of correlations was also calculated using permutation test. The label of each trial was shuffled and 1000 iterations of the whole decoding procedure and correlation analysis was performed to get the null distribution of partial Pearson correlation r. P value was calculated by the number of times that the r in the set of 1000 permutation tests outperformed the real r divided by 1000. P values were corrected for multiple comparisons using FDR.

### 2.11 Whole brain decoding map and conjunction analysis

Trying to get the full picture of brain regions sensitive to distance/magnitude, the SVR decoding models across the whole-brain was run, interpreting the model weight of voxels as a decoding map. To select most important voxels, bootstrap tests with 1,000 samples for each subject were performed (resampling with replacement)(Kohoutová et al., 2022). For every voxel in each individual, uncorrected P values were derived based on the sampling distribution. These P values were obtained by converting z-scores to probabilities using the mean and standard deviation of sampling distribution. Subsequently, negative logarithmic P values (mean (−log(P))) was computed and voxels corresponding to the top 10% of these mean (−log(P)) values were identified. These selected voxels constituted our predictive map for distance and magnitude.

The binarized predictive map of distance and magnitude were utilized as a mask. By overlaying this mask, a conjunction map was generated. The conjunction map revealed common brain regions that play significant roles in representing both distance and magnitude.

## 3. Result

### 3.1 Decoding results of number distance and magnitude

To test whether participants formed a distance representation (the distance between the first and second dot pattern) during the dot magnitude comparison task, support vector regression (SVR) was employed to decode number distance. The intra-parietal sulcus (IPS) and the hippocampus (HIPP) were chosen as regions of interests, and fine-grained information was extracted from multivoxel pattern. The distance of each trial was defined as the absolute value of the difference between the first and second dot, treated as continuous variables, and regression was used to detect gradual changes in multivoxel patterns as a function of numerical distance.

The theoretically expected performance in noninformative data corresponds to 0, so the significance level was generated by group level t test against 0. Permutation test was then conducted to further confirm the reliability of these results. The results showed that patterns in IPS were able to predict the number distance (Fig:3B) : IPS (*z_r_* = 0.30 ± 0.02 (mean ± s.e.m.) with 5-fold cross-validation, *p_perm_* < 0.001), RIPS (*z_r_* = 0.29 ± 0.02, *p_perm_* < 0.001), LIPS (*z_r_* = 0.23 ± 0.03, *p_perm_* < 0.001). However, patterns in hippocampus were not able to predict the distance (Supplementary Fig.2B): HIPP (*z_r_* = 0.06 ± 0.02, *p_perm_* = 0.099), RIPS (*z_r_* = 0.05 ± 0.03, *p_perm_* = 0.129), LIPS (*z_r_* = 0.06 ± 0.03, *p_perm_* = 0.099). This indicated that the distance information was detected in IPS when subjects performed the number comparison task. RIPS exhibited greater precision in distance decoding compared to LIPS (Fig.3D, paired samples t-test: *t*=2.728, *p*=0.011, *p_corrected_=*0.039).

Similar to distance decoding, parallel analysis was conducted to decode number magnitude (the magnitude of the second dot pattern in the comparison task) in IPS ROIs. The decoding results were shown in Fig.3C. Patterns in all ROIs were able to predict the number magnitude: IPS (*z_r_*=0.25±0.02 (mean ± s.e.m.) with 5-fold cross-validation, *p_perm_*<0.001), RIPS (*z_r_* = 0.21±0.03, *p_perm_*<0.001),LIPS (*z_r_* =0.25±0.03, *p_perm_*<0.001). The results imply that the subjects access the magnitude representation during the number comparison task as well. There was no differences in magnitude decoding performance between LIPS and RIPS, but we found that distance decoding performance was higher than magnitude in RIPS (Fig.3D, paired samples t-test: RIPS vs LIPS(mag), *t*=1.376, *p*=0.180, *p_corrected_=*0.240; mag vs dis(RIPS), *t*=2.476, *p*=0.020, *p_corrected_=*0.039;mag vs dis(LIPS),*t*=0.026, *p*=0.795, *p_corrected_=*0.795).

### 3.2 Representational similarity analyses

To provide additional evidence on effect of distance and magnitude representation on the task, the alignment of theoretical model of tuning curves overlapping (Lyons, Ansari, et al., 2015) with the observed similarity of neural patterns or behavior, distance or magnitude were tested.

#### 3.2.1 Behavioral Patterns of Number Distance and Magnitude Representations

Firstly, it was examined whether the overlapping model of distance or magnitude could account for the behavioral RDM. The trial-by-trial Euclidean distance of RT was used to describe behavioral dissimilarity in-between. The overlap of the tuning curves for the two consecutive numbers within each trial was used to generate a graph representing the magnitude overlap curve (Fig 2A, where the shaded area shows the overlap proportion of the tuning curves for the two consecutive number magnitudes). Each curve reflects the overlap of response elicited by a pair of numbers in a trial, and the overlaps of these curves (‘overlap of overlap’) indicate the similarities between trials (Fig 2B). So the “overlap” of overlapping tuning curve was then taken as the magnitude overlap to see the relationship of it with the dissimilarities of RT between trials (Fig 4A). Note that, the trial-by-trial model RDM was calculated by 1-overlap. Simultaneously, a distance model was applied to establish tuning curves representing the distance between the two numbers within each trial, and the overlap of these distance tuning curves (referred to as "distance overlap") was used to describe trial-to-trial similarity (Fig 2C, Fig 4A).

**Figure 2.**
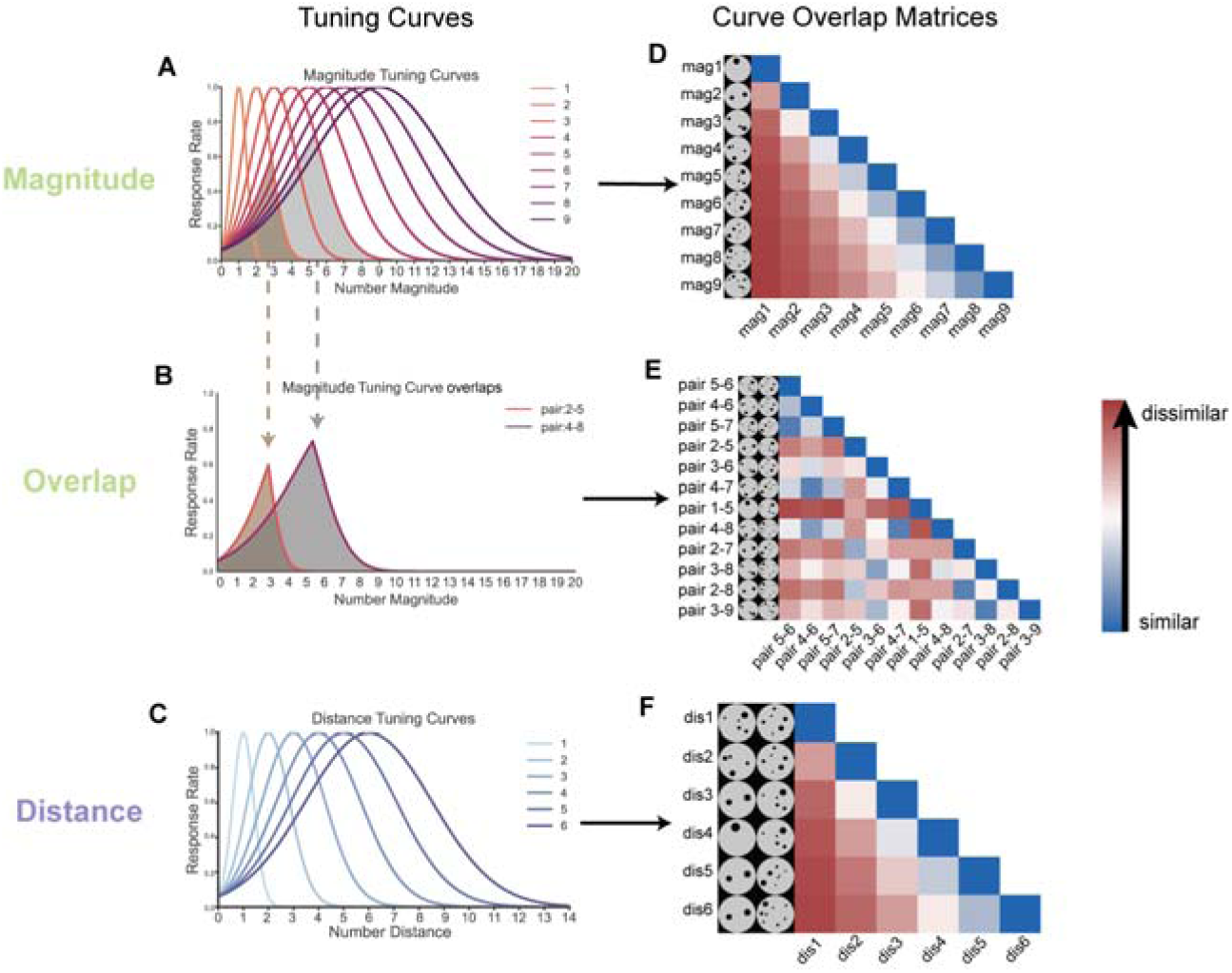
Generating Curve Overlap Matrices. Note: Simulated neuronal tuning curves for number magnitude (A), distance (C) and overlap curve for magnitude (B); Gaussian functions were computed with width linearly increasing with number magnitude or distance, consistent with human behavioral data. (D-F) For each pair of tuning curves (overlap curves), these shows 1-the proportion of those curves that overlap with one another. Note when the overlap increases, the dissimilarity between two curves decreases.

**Figure 3.**
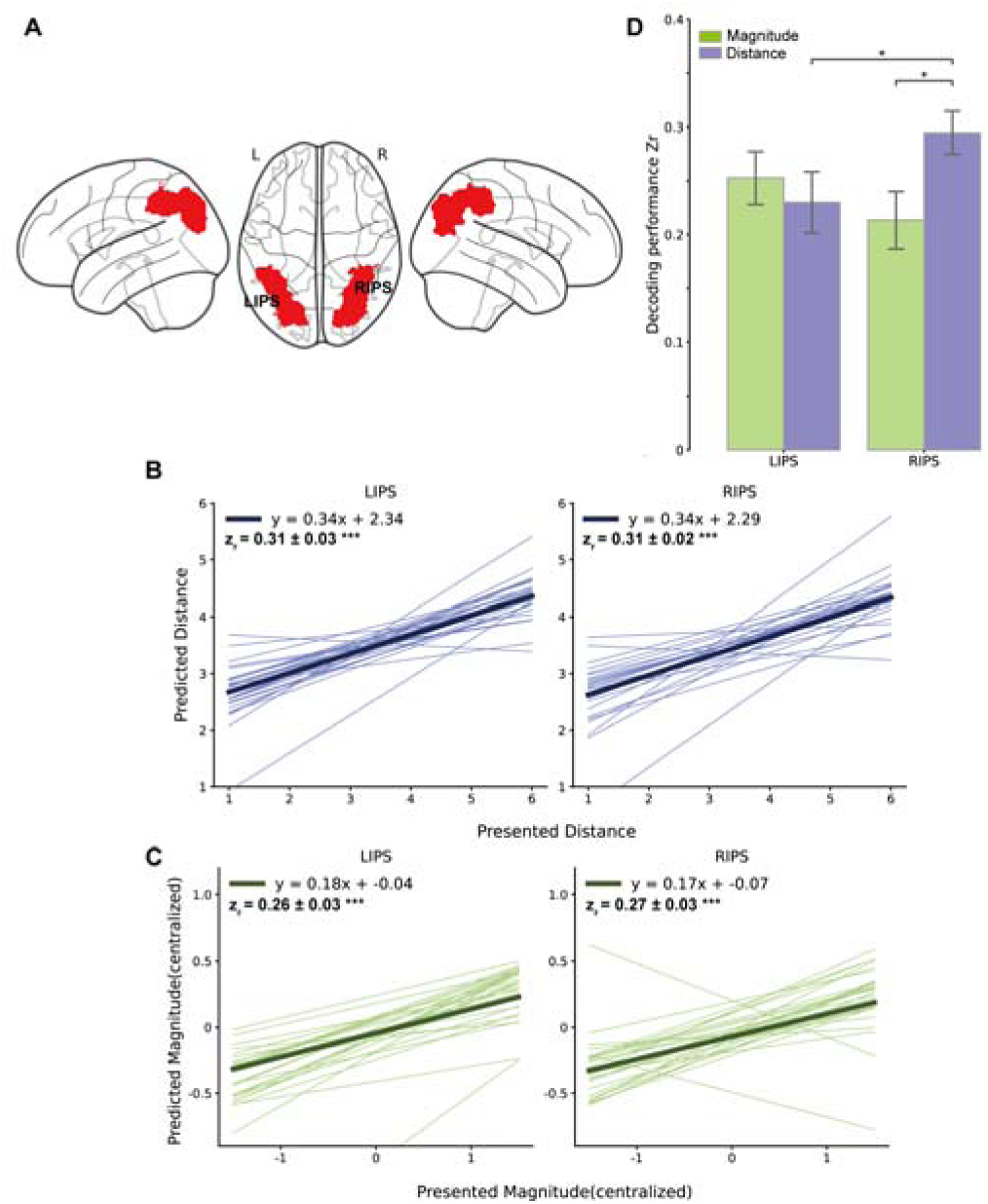
ROIs and SVR Decoding Results. Note: (A) Regions of interest ware defined according to the Julich Brain Atlas. (B-C) The magnitude and distance (the absolute value of the difference between two numbers) were significantly decoded from all three ROIs. Decoding performance was measured using Pearson’s correlations between the distance (B) or magnitude (C) of presented number and their predictions obtained using SVR. Each thin line represents an individual subject, the thick line represents the population average. SVR model were tested in 5-Fold cross validation. *z_r_*was fisher z transformed *r* and averaged across subject(mean ± s.e.m.). (D) Differences in decoding performance among ROIs. Fisher Z transformed from Pearson’s *r* were averaged across subjects within each region of interest. Error bars reflect s.e.m.(\**P_corrected_*<0.05, \*\**P_corrected_*<0.01,***,*P_corrected_*<0.001).

Results indicated that both models, to some extent, captured behavioral RT geometry (Raw effect: magnitude: *z*=0.023, *p*=0.010; distance: *z*=0.070, *p*<0.001; Preliminary effect: magnitude: *z*=0.023, *p*=0.010; distance: *z*=0.070, *p*<0.001, Fig4B). However, when mutually controlling for each other, only the unique effect of the distance model remained significant (Unique effect: magnitude: *z*=0.002, *p*=0.42; distance: *z*=0.066, *p*<0.001). This result suggests that the representation of distance between presented numbers within a trial plays a crucial role in explaining behavioral similarity than the overlaps of magnitude tuning curves.

#### 3.2.2 Neural Patterns of Number Distance and Magnitude Representations

It was assumed that the correlation between the distributed patterns of activity for two trials is indeed predicted by the extent to which their tuning curves overlap. Consequently, two types of trial-by-trial Representational Dissimilarity Matrices (RDMs) for magnitude and distance were defined based on tuning curve overlap model. To examine the magnitude representation, magnitude RDM was defined to reflect the overlap of the tuning curves for the second number of each trial (referred to as "magnitude of number 2 overlap"). Concurrently, the same RDM for distance representation as employed in our behavioral RSA analysis was used. These RDMs were correlated with neural RDM (Fig4C), and this correlation was calculated using only across-run trial pairs to control for Type I error(Mumford et al., 2014). Initially, Spearman correlation was used to test the raw effect. Then, visual and response RDMs were included as confound RDMs, and partial Spearman correlation was calculated to test the preliminary effect. Subsequently, the unique effect was introduced by adding another computational model as a confound RDM.

The results in IPS replicated the findings from Lyons et al. (2015) for magnitude representation, demonstrating a correlation between the overlaps of magnitude tuning curves and neural pattern similarity. Additionally, the overlaps of distance tuning curves exhibited a similar pattern (Raw effect: magnitude: *z*=0.006, *p*=0.042; distance: *z*=0.011, *p*=0.042). These indicated that both magnitude and distance models captured neural similarity to some extent. However, after controlling for visual and response RDMs, the significance of the magnitude model diminished, while the distance model significantly correlated with neural similarity (Fig.4D, Preliminary effect: magnitude: *z*=0.004, *p*=0.101, distance: *z*=0.011, *p*=0.042; Unique effect: magnitude: *z*=0.004, *p*=0.101; distance: *z*=0.011, *p*=0.042. supplementary table 2).

**Figure 4.**
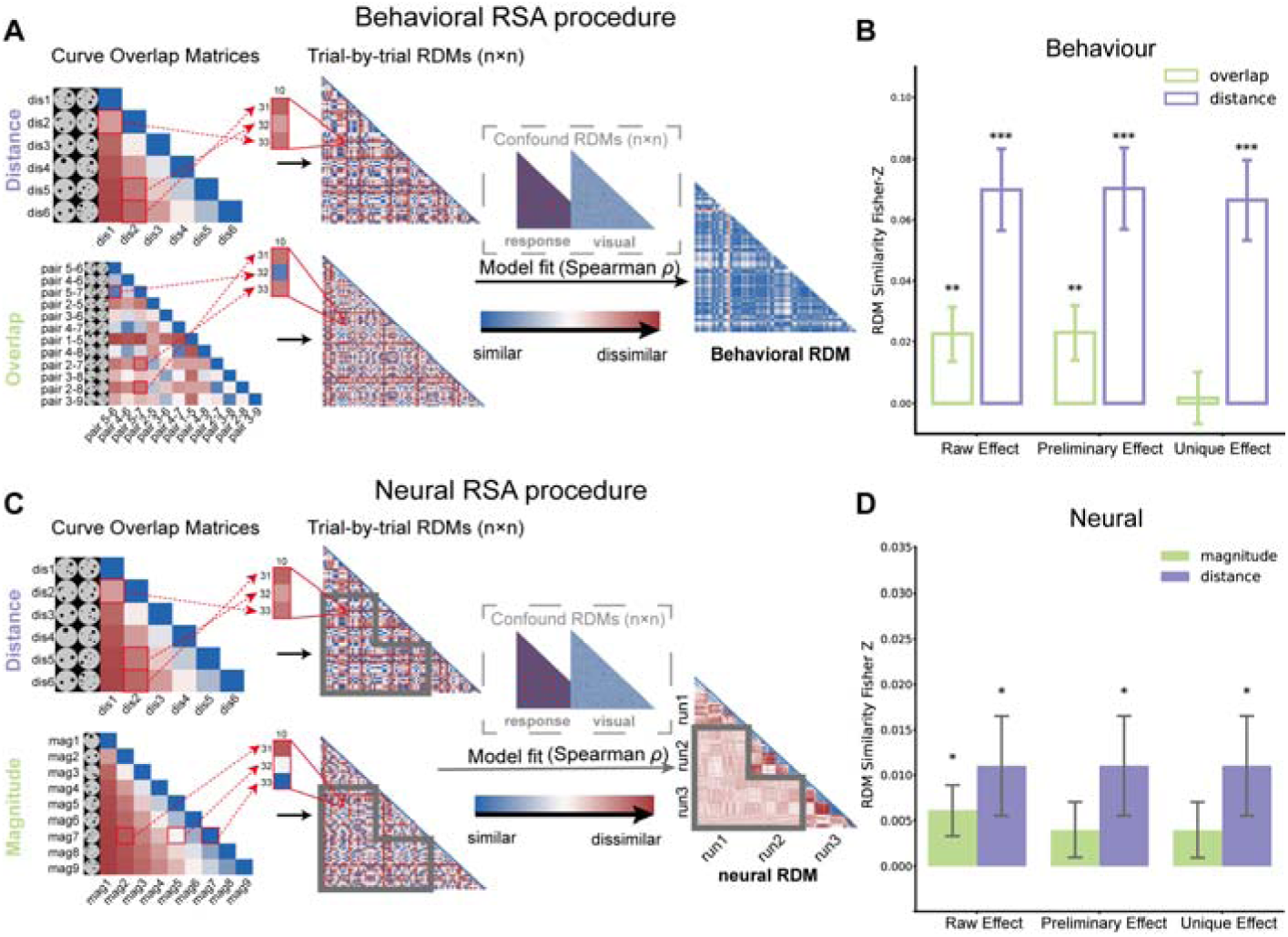
Representational Similarity Analysis (RSA) Procedure and Results. Note: This diagram outlines the RSA procedure connecting computational models with brain activity and behavioral data. The left column shows the Curve Overlap Matrices for distance **(A, C)**, magnitude **(C)** and overlap **(A)** which was generated from tuning curve overlap; The middle column displays trial-by-trial Representational Dissimilarity Matrices (RDMs) generated from Curve Overlap Matrices according to each trial’s category. The right column exhibits trial-by-trial behavioral RDM which was defined by the Euclid distance of RT **(A)** and the neural RDM **(C)** calculated by Pearson correlation. Notably, only between-run trials were compared to neural RDM (indicated with grey frames, C), while behavior employed full trials. Spearman correlation is employed to associate model RDMs with behavior or neural RDMs. **(B)** Both the distance and overlap RDM significantly explain the behavioral differences in reaction times (RT) across trials, but only distance unique effect is significant. **(D)** Both distance RDM and magnitude RDM could significantly explain neural representational geometry in IPS, but only distance preliminary and unique effect are significant. Bar graph shows Fisher Z transformed Spearman correlation (partial) coefficient. Group means and standard errors of the similarity are indicated by the bar plots with error bars. Significance of similarity is tested using one-sample t-test and indicated by black asterisk: \**p* < .05, \*\**p*<0.01,\*\*\**p*<0.001, FDR-corrected.

The results in hippocampus indicated that distance models cannot captured neural similarity in hippocampus (Raw effect: BHIPP: *z*=0.006, *p*=0.234, LHIPP: *z*=0.007, *p*=0.159; RHIPP: *z*=0.003, *p*=0.312, Supplementary Fig.2C).

### 3.3 Neural distance representation precision related to individual difference of numerical acuity and math performance

After confirming the existence of distance representation through both decoding and RSA analyses, the aim was to define a neural index that describes the precision of representation during a number comparison task. Notably, a higher *z_r,_* between the SVR decoder’s prediction and the presented one for a participant indicated that the neural patterns in the IPS represented distance more accurately. Thus, *z_r_* was defined as each individual’s representation precision (RP). It was assumed that individuals would demonstrate greater numerical acuity and better math performance when they have more accurate distance representation.

Consistent with the hypothesis, it was found that distance RP was negatively correlated with the Weber Fraction score (Table.1), which was assessed by psychophysical modelling of performance on the number comparison task. A higher Weber fraction score corresponds to poorer ANS acuity. Correlation: IPS (*r*=0.-560, *p*=0.002, *p_perm_*=0.005). The partial correlation results are shown in Fig.5A: IPS (*r*=0.-551, *p*=0.007, *p_perm_*=0.009). This indicates that individuals with higher neural representation precision of distance have better ANS acuity.

**Table 1.** Distance RP Partial Correlations with Arithmetic Performance and Weber Fraction.

|  | Region | Correlation |  | Partial correlation |  |
| --- | --- | --- | --- | --- | --- |
| | | Pearson's $r$ | $p$-corrected | Pearson's $r$ | $p$-corrected |
| **Arithmetic** | LIPS | 0.534713 | 0.003 | 0.456889 | 0.025 |
|  | RIPS | 0.509626 | 0.004 | 0.388181 | 0.025 |
|  | BIPS | 0.500787 | 0.004 | 0.383701 | 0.025 |
| **Weber fraction** | LIPS | -0.483441 | 0.005 | -0.412170 | 0.014 |
|  | RIPS | -0.521000 | 0.005 | -0.481244 | 0.0105 |
|  | BIPS | -0.559569 | 0.005 | -0.551130 | 0.009 |

**Figure 5.**
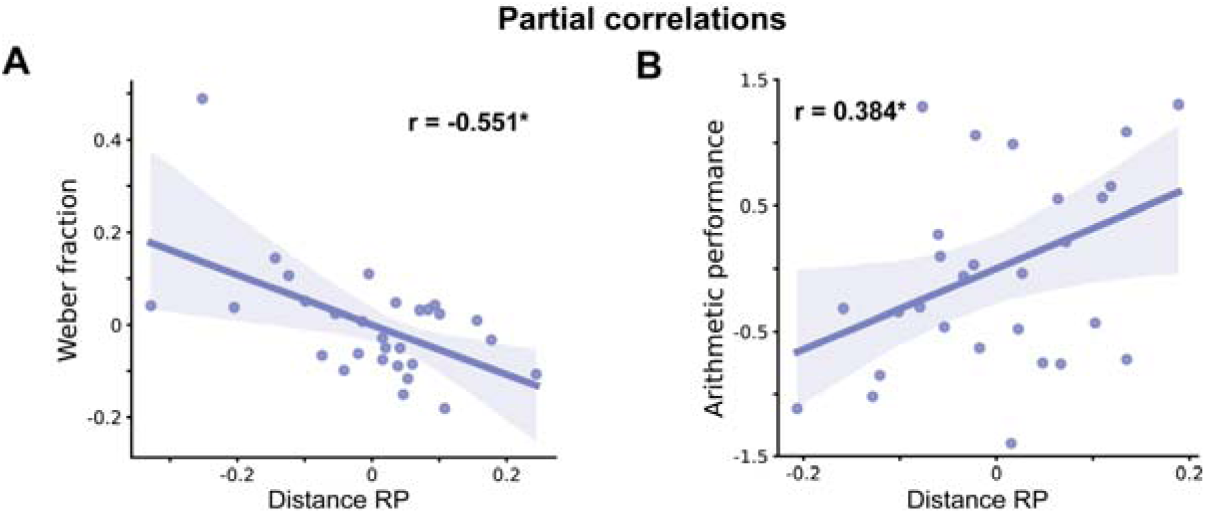
Partial Correlations. Note: The SVR decoding performance of distance *Z_r_*for each subject was defined as representation precision (RP). **(A)** Distance RP (decoding performance *Z_r_*) in the IPS was significantly correlated with the Weber fraction (estimated in number comparison task, with lower value indicating better performance) even after controlling for RT. **(B)** Distance RP also showed a significant association with arithmetic performance after controlling for the RT and accuracy of dot comparison task. Significance of partial correlation is indicated by black asterisk: \**p* < .05, *p* value was generated from permutation test, FDR-corrected.

The correlations between the participants’ distance RP and their symbolic arithmetic performance were then examined (Table.1). Positive correlations were observed between RP and arithmetic performance in IPS (*r*=0.501, *p_perm_*=0.004), even after controlling for RT and accuracy of the dot comparison task: IPS (partial *r* = 0.384, *p_perm_*=0.025, Fig.5B). These results suggest that individuals who can represent distance more precisely tend to perform better in symbolic arithmetic task.

The correlations between the magnitude RP of the participants and their Weber fraction and symbolic arithmetic performance were also tested. No correlation was found between magnitude representational precision (RP) and the Weber Fraction score. The negative correlations between RP and arithmetic performance in IPS were only marginal significant (*r*=-0.343, *p_perm_*=0.084), but when RT and accuracy were taken into account, this association was no longer significant (IPS: *r*= -0.320, *p_perm_* = 0.149, Supplementary Fig.1,Table.3).

### 3.4 Whole brain maps for distance or magnitude decoding

To further detect distance sensitive brain regions other than IPS, the SVR models were applied to each participant’s whole-brain multivoxel pattern, creating 29 individualized distance-decoding maps. Important features for distance decoding were identified using bootstrap tests with 1000 iterations for each individual’s map. With the P values from bootstrap tests, the mean (-log(P)) values were computed for the whole brain across all individualized maps, and the top 10% voxels were selected, with a threshold of mean (-log(P)) = 1.649. The selected voxels are displayed in Suppl. Fig.3A and Suppl. Table 4. The distance sensitive map involved widespread region in the cortex, including visual cortex, IPS, angular gyrus, precuneus, motor cortex, prefrontal cortex, insula, medial temporal gyrus, cingulate cortex. The same procedure was applied to find important magnitude-sensitive brain regions, selecting the top 10% voxels, with the threshold being the mean (-log(P)) = 1.827. The selected voxels are displayed in Suppl. Fig.3B and Suppl. Table 4. The magnitude predictive map involved widespread region in the cortex.

Furthermore, an conjunction analysis was performed to identify the common region of the two predictive maps. The results showed that neural signals in bilateral intraparietal sulcus, left temporal-occipital area and lingual gyrus play an important role in representing both magnitude and distance (Suppl. Fig.3C; Suppl. Table 4).

## 4. Discussion

Generally, the present study has revealed the significant influence of distance representation in the magnitude comparison task and the importance of its precision in both the task and mathematical abilities. Through neural activation patterns in the IPS, the study demonstrated that distance information was decoded between presented pair of dots. A correlation was found between the precision of distance presentation, as indicated by decoding accuracy, and both the Weber fraction and arithmetic performance.

Using a neural decoding method, it was discovered that the multivoxel activation pattern in the IPS following the presentation of the second stimulus differentiated the distance between the two magnitudes compared, thus successfully predicting those distance. These results firstly revealed the importance of distance representation in magnitude comparison task, complementing previous investigations of magnitude representation. While this study replicated the successful decoding of magnitude as reported in previous findings, it was also found that overlaps in neural tuning curves accounted for reaction time RDM and neural RDM, in models of both magnitude and distance. The successful decoding of magnitude and the replication of tuning curve overlapping model of magnitude (Lyons, Ansari, & Beilock, 2015) confirmed the access of magnitude representation during magnitude comparison task. However, when mutually controlling for each other, only the distance model survived. The distance model could still explain the neural RDM even when controlling for visual and response RDMs, whereas the magnitude model could not. These results suggest that when people compare two magnitudes, they represent each magnitude and compare them based on these representations. The overlapping of the neural curve initiated by each magnitude influences behavioral performance, with more overlap leading to increased difficulty. Moreover, not only is magnitude representation activated during comparison, but individuals also “measure” the difference between the two stimuli with a “ruler”, referred to distance representation. Distance representation integrates the relational information between stimuli, making it more complex and demanding than representing only one of them. Consequently, distance representation is more critical than magnitude representation in magnitude comparison. Considering that previous studies have claimed distance representation in multidimensional physical (Howard et al., 2014; Morgan, Macevoy, Aguirre, & Epstein, 2011) or abstract semantic spaces. Researchers have discovered that both univariate activation and multivariate pattern encoded Euclidean distance between individual positions in object, word (Viganò & Piazza, 2020; Viganò, Rubino, Soccio, Buiatti, & Piazza, 2021), and concept spaces (Theves, Fernandez, & Doeller, 2019). These findings indicate that representing relational information-specifically distance – is crucial to identify the relationships between items. Such evidence inferred that distance representation may serves as a domain-general foundation for human cognition and learning. Consequently, it would be highly promising to further investigate distance representation across various domain in future research.

Although the brain location of the distance representation is not the primary focus of the current study, it is noteworthy that the previous studies on navigation in map-like space has found that distance is encoded mainly in the hippocampus or entorhinal cortex, rather than in the parietal or intraparietal sulcus. One possible reason is that the parietal was not selected as a region of interest in studies related to cognitive space, leading to its neglect to some extent. Alternatively, different brain regions may be sensitive to one-dimensional or multi-dimensional spaces. Additionally, the hippocampus might be involved during the initial phases of map construction with the parietal cortex assisting in constructing the map being too weak to detect initially. As the internal space matures, distance representation may switch from subcortical hippocampus to cortical parietal cortex, as described by the complimentary learning system(McClelland, McNaughton, & O’Reilly, 1995). Those hypotheses warrant further investigation. And the widespread distinct brain areas involved in distance and magnitude representation other than the IPS revealed by whole brain analysis suggest a brain network perspectives for future study. The common decoding area in bilateral IPS confirm again the numerical-related and math related function in IPS.

Our study also found that the precision of distance representation is crucial in basic numerical processing and math cognition. The present study showed a positive correlation between the precision of distance representation and both number acuity and arithmetic performance, but not magnitude representation. The Weber fraction, indicating number acuity, describes the limit of the finest numerical changes an individual can reliably detect. Thus it is reasonable that a person with higher acuity can more precisely determine the difference between two numbers with more accurate distance representation. While the accurate location of magnitude representation facilitates precise distance representation, it is not fully conditional. Our result also found that the magnitude representation precision was not directly correlated with the Weber fraction. This partially explains the controversial results regarding the relationship between magnitude processing and mathematical abilities. The precise location of the two stimuli is only the first step in the comparison; accurately detecting the distance between them has a more direct impact. It can be assumed that if a person can measure the distance between different magnitudes, they might be more sensitive to different numbers and their numerical relationship. Consequently, they would perform better or more efficiently during arithmetic tasks or when exploring the relationship between given conditions and goals in mathematical problem-solving. This is why the present study not only showed a positive correlation between the accuracy of distance decoding and the number acuity but also arithmetic performance. The ability to accurately detect distance representation may also explain why some individuals are better at identifying relationships between various variables, and demonstrating talents in mathematics or problem-solving.

Generally, the current study is the first to highlight the critical role of distance representation in magnitude comparison tasks and its broader implications for mathematical abilities. Our findings demonstrate that precise distance representation, rather than magnitude representation alone, significantly predicts task performance and arithmetic proficiency. However, this study on distance representation is quite preliminary. More concrete evidence is needed to draw a clearer picture of distance representation. Future research should further explore the cognitive components and mechanisms underlying the distance representation, including the involvement of possible distinct neural regions in representing different dimensional space, as well as the attribution of distance representation across different domains. Exploring these areas will provide deeper insights into the cognitive and neural foundations of relational processing, potentially informing strategies to enhance cognitive abilities across various domains, beyond just numerical skills.

## Contribution

Conceived and designed the experiments: HZ. Investigation: HZ JY JL. Formal analysis: HZ WQ YY JL JY. Methodology and Software: WQ JX SQ. Writing-original draft: HZ WQ. Writing-review and editing: HZ WQ SQ. Supervision and Project administration: HZ and SQ.

## Acknowledgment

The authors thank Hao Lu, Yi Feng and Hailian Hu for their assistance for data collection.

This study was supported by National Natural Science Foundation of China(62077010); the STI 2030—Major Projects (2021ZD0200500); Open Research Fund of the State Key Laboratory of Cognitive Neuroscience and Learning (CNLYB2103).

## Data Sharing and Data Availability

The functional imaging data examples and code for decoding was available in https://github.com/qwhying/NRPD-predict-arithmetic/tree/main.

