## Supplementary materials for "Neural Representation Precision of Distance Predicts Arithmetic Performance"

**Supplementary Table 1**

*Arithmetic Fluency Test performance of primary school students dataset*

| Grade | *N* | | *M* | *SD* |
| --- | --- | --- | --- | --- |
|  | Boys | Girls |  |  |
| 1 | 180 | 155 | 0.73 | 0.26 |
| 2 | 170 | 161 | 0.89 | 0.27 |
| 3 | 168 | 156 | 2.14 | 0.59 |
| 4 | 183 | 148 | 2.44 | 0.64 |
| 5 | 154 | 170 | 2.78 | 0.58 |
| 6 | 146 | 164 | 2.97 | 0.62 |

**Supplementary Table 2**

*Neural RSA Result*

|  | **Magnitude Model** | | | **Distance Model** | | |
| --- | --- | --- | --- | --- | --- | --- |
|  | *z* | *p* | *p_corrected_* | *z* | *p* | *p_corrected_* |
| **Raw effect** | 0.006104 | 0.018008 | 0.042 | 0.011008 | 0.027940 | 0.042 |
| **Preliminary effect** | 0.003989 | 0.099372 | 0.101 | 0.011028 | 0.027743 | 0.042 |
| **Unique effect** | 0.003971 | 0.100596 | 0.101 | 0.011021 | 0.027851 | 0.042 |

**Supplementary Table 3**

*Magnitude RP Partial Correlation with Weber Fraction and Arithmetic Performance*

|  |  | **Correlation** | | **Partial correlation** | |
| --- | --- | --- | --- | --- | --- |
|  | region | Pearson’s *r* | *p*_corrected_ | Pearson’s *r* | *p*_corrected_ |
| **Arithmetic** | LIPS | -0.259952 | 0.107 | -0.251205 | 0.149 |
|  | RIPS | -0.323350 | 0.084 | -0.253877 | 0.149 |
|  | BIPS | -0.342540 | 0.084 | -0.320495 | 0.149 |
| **Weber fraction** | LIPS | 0.197165 | 0.117 | 0.165360 | 0.156 |
|  | RIPS | 0.358350 | 0.072 | 0.287661 | 0.144 |
|  | BIPS | 0.252852 | 0.117 | 0.219142 | 0.144 |

**Supplementary Table 4**

*Brain regions of whole brain analysis*

| Brain regions | Cluster size  (voxels) | Hem | MNI Coordinates（mm） | | | Peak Z |
| --- | --- | --- | --- | --- | --- | --- |
|  |  |  | X | Y | Z |  |
| Distance |  |  |  |  |  |  |
| Precuneus | 2104 | R | 6 | -60 | 62 | 2.23 |
| Intraparietal sulcus | 1638 | L | -45 | -63 | 21 | 2.38 |
| Angular | 364 | R | 40 | -65 | 39 | 2.31 |
| Supra Marginal  gyrus | 1115 | L | -20 | -78 | 34 | 2.19 |
| Superior frontal gyrus | 888 | R | 2 | 25 | 40 | 2.40 |
| Middle frontal gyrus | 384 | L | -33 | 23 | 44 | 2.25 |
| Middle temporal gyrus | 308 | R | 53 | -41 | -12 | 2.24 |
| Superior temporal gyrus | 325 | L | -57 | -18 | 6 | 2.23 |
| Superior occipital gyrus | 251 | L | -18 | -69 | 15 | 2.46 |
| Precentral gyrus | 257 | R | 45 | -21 | 47 | 2.19 |
| Insula | 857 | L | -38 | 20 | -6 | 2.74 |
| Anterior cingulate | 266 | R | 1 | 43 | 11 | 2.23 |
| Middle cingulate | 289 | R | 1 | -26 | 46 | 2.24 |
| Hippocampus | 284 | L | -25 | -37 | -8 | 2.35 |
| Calcarine | 1347 | L | -3 | -89 | 9 | 2.54 |
| Cerebellum | 221 | L | -4 | -39 | -27 | 2.32 |
| Cerebellum | 464 | R | 22 | -77 | -21 | 2.58 |
| Magnitude |  |  |  |  |  |  |
| Fusiform gyrus | 4874 | L | -17 | -67 | 16 | 2.93 |
| Superior occipital | 344 | L | -20 | -78 | 34 | 2.19 |
| Intraparietal sulcus | 1172 | R | 44 | -55 | 37 | 2.64 |
| Superior frontal gyrus | 374 | L | -24 | 50 | 28 | 2.65 |
| Inferior frontal gyrus | 279 | L | -50 | 22 | 19 | 2.78 |
| Inferior frontal gyrus | 264 | R | 51 | 12 | 14 | 2.41 |
| Middle frontal gyrus | 342 | R | 28 | 38 | 38 | 2.90 |
| Precentral gyrus | 719 | L | -35 | 0 | 51 | 2.51 |
| Superior temporal gyrus | 230 | L | -60 | -37 | 14 | 2.63 |
| Middle temporal gyrus | 439 | L | -60 | -22 | -7 | 3.12 |
| Anterior cingulate | 477 | L | -4 | 46 | 0 | 2.48 |
| Middle cingulate | 647 | L | -1 | -15 | 45 | 2.82 |
| Para-Hippocampal | 203 | L | -32 | -28 | -16 | 2.53 |
| Insula | 299 | R | 47 | -10 | 14 | 2.76 |
| Conjunction |  |  |  |  |  |  |
| Intraparietal sulcus | 121 | L | -36 | -71 | 41 |  |
| Intraparietal sulcus | 57 | R | 50 | -44 | 39 |  |
| Superior occipital gyrus | 73 | L | -20 | -72 | 25 |  |
| Middle temporal gyrus | 71 | L | -45 | -68 | 21 |  |
| Lingual gyrus | 74 | L | -17 | -68 | 6 |  |

**Supplementary Figure 1**

*Partial Correlations of Magnitude RP.*

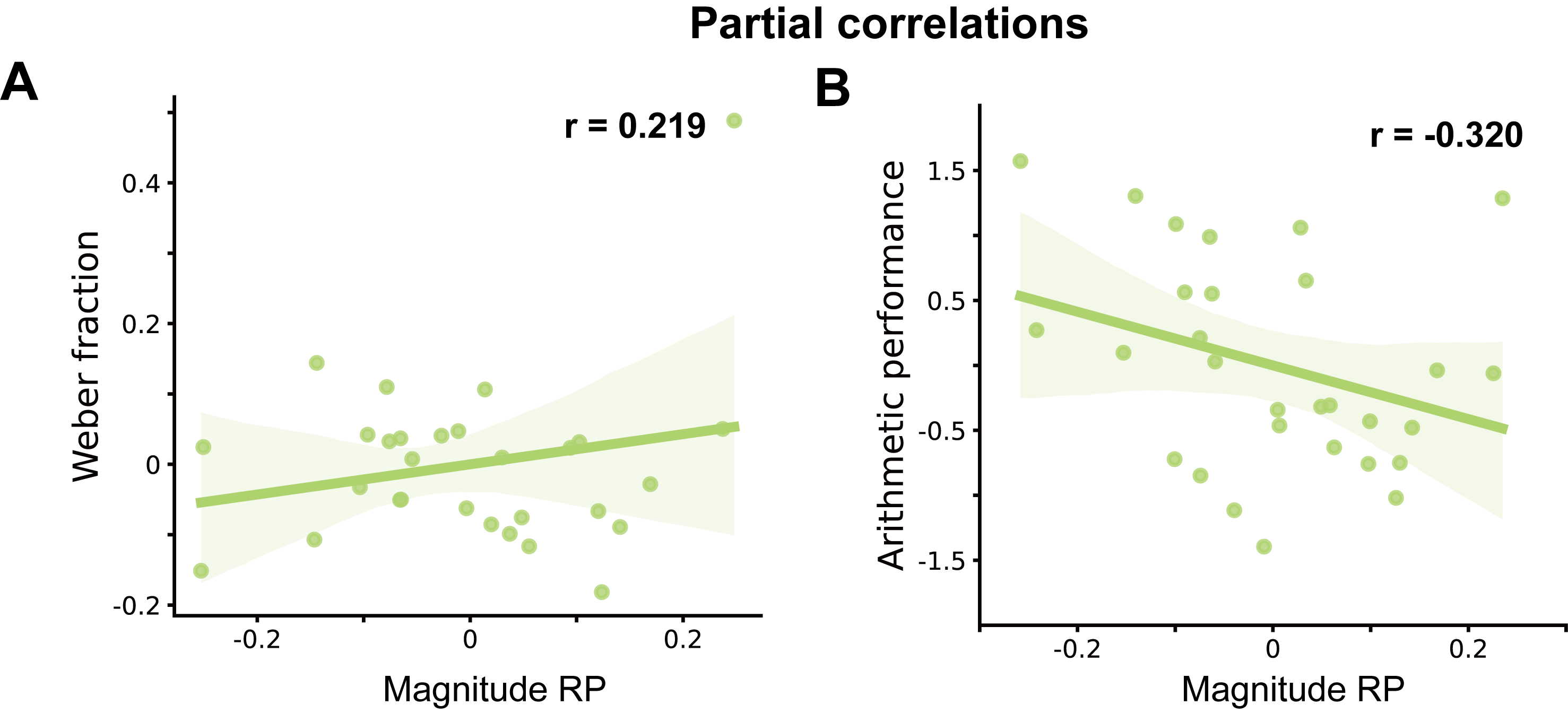

**(A)** Magnitude RP (decoding performance *Z_r_*) in the IPS was not significantly correlated with the Weber fraction (estimated in number comparison task, with lower value indicating better performance) after controlling for the RT. **(B)** Magnitude RP also did not correlate with arithmetic performance after controlling for the RT and accuracy of dot comparison task. Significance of partial correlation is indicated by black asterisk: **p* < .05, *p* value was generated from permutation test, FDR-corrected.

**Supplementary Figure 2**

*Decoding in Hippocampus.*

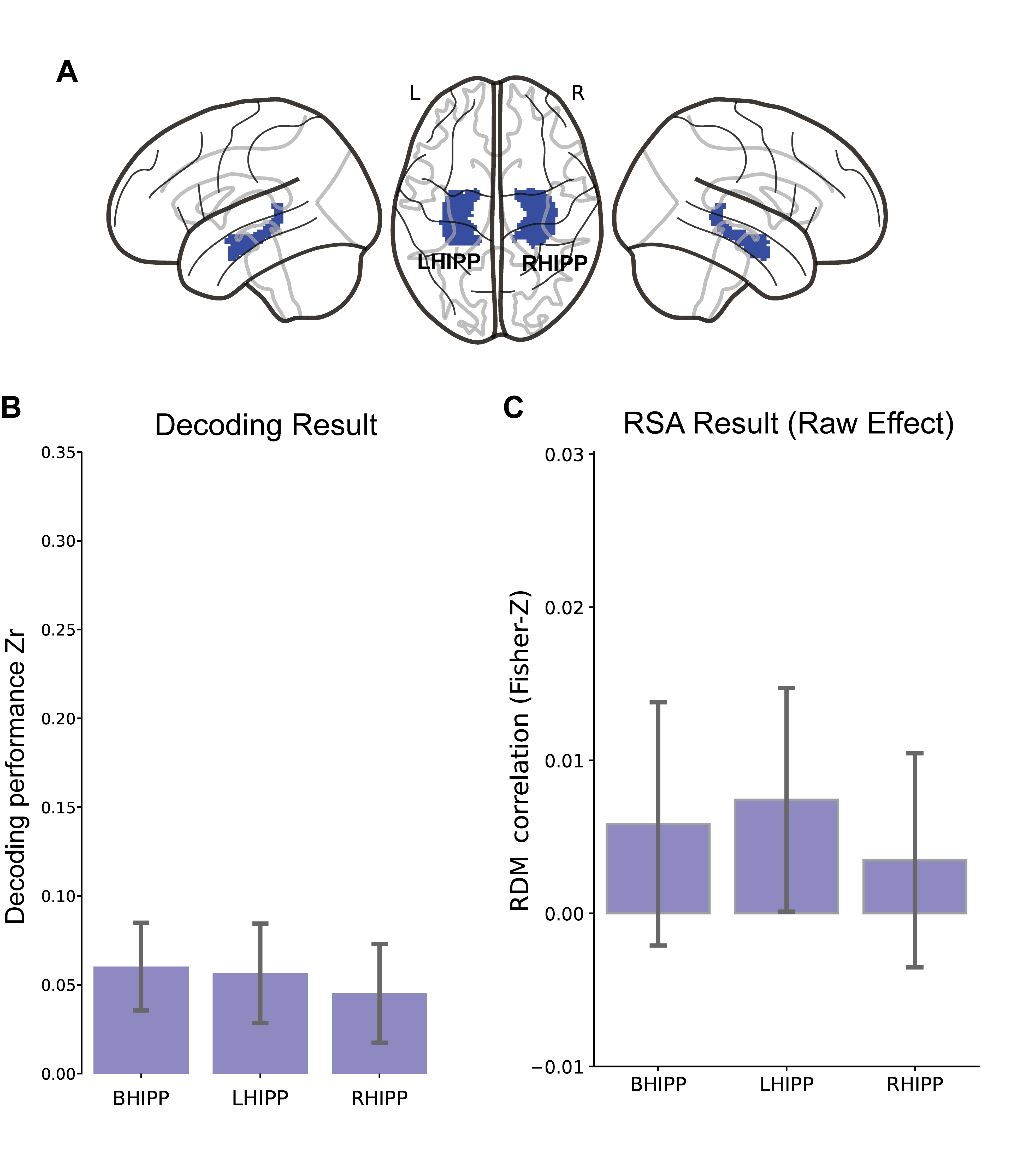

**(A)** Regions of interest ware defined from the WFU pickAtlas **(B)** Decoding performance among ROIs, results showed that patterns in the hippocampus were not able to predict the number distance. **(C)** Distance models cannot capture neural similarity in the hippocampus.

**Supplementary Figure 3**

*Whole Brain Decoding Maps of Magnitude and Distance*

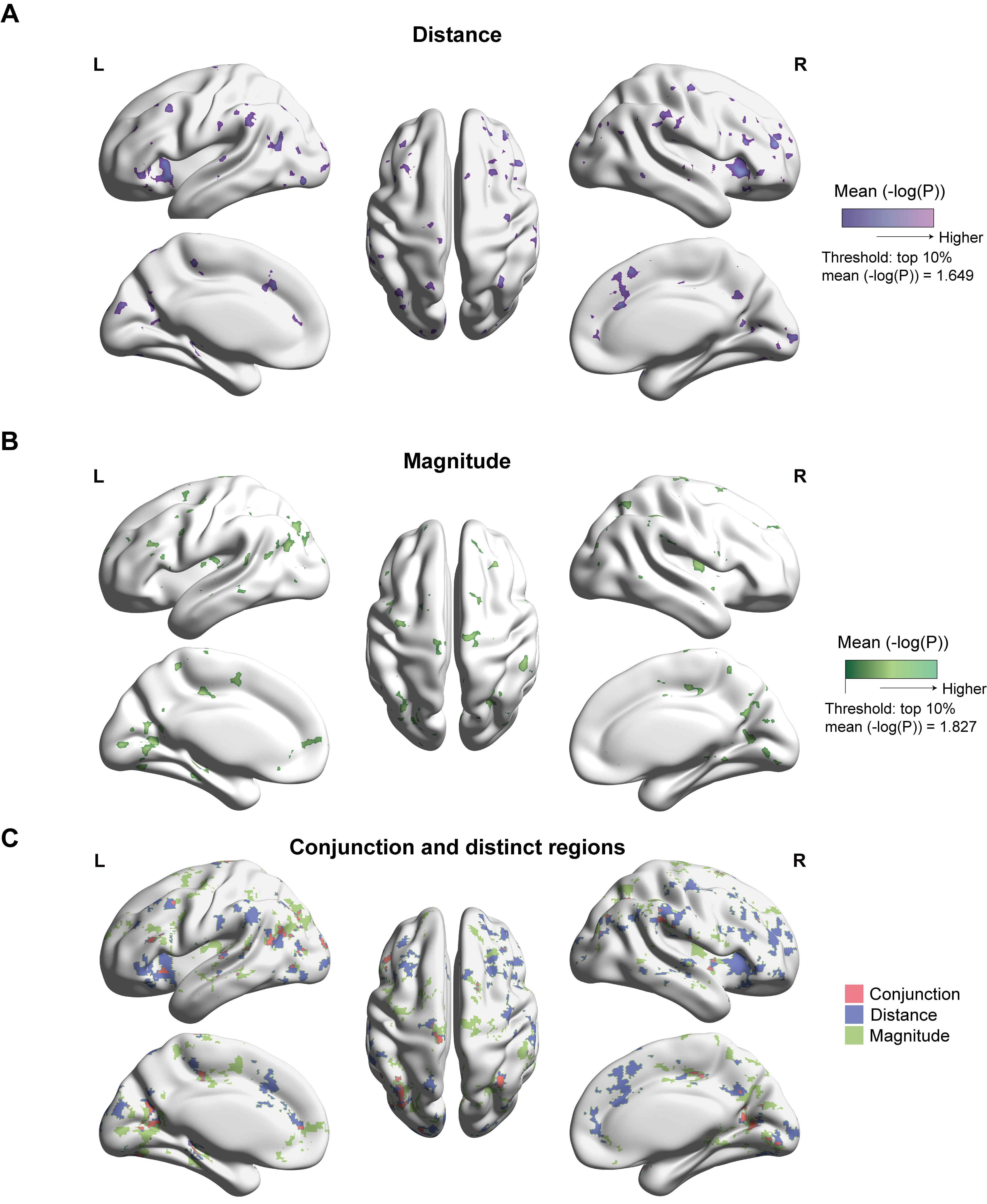

Note: (A) Selected voxels important for magnitude prediction. The importance was operationalized in terms of mean (−log(*P*)), where the two-tailed *P* values were calculated from the bootstrap tests performed in each individualized decoding map. The selected voxels had mean (−log(*P*)) values higher than the top 10% mean (−log(*P*)) value (=1.827). (B) Selected voxels important for distance prediction. The selected voxels had mean (−log(*P*)) values higher than the top 10% mean (−log(*P*)) value (=1.649).(C) Conjunction and unique area of distance and magnitude predictive maps.
